# Multiple introductions of Zika virus into the United States revealed through genomic epidemiology

**DOI:** 10.1101/104794

**Authors:** Nathan D Grubaugh, Jason T Ladner, Moritz UG Kraemer, Gytis Dudas, Amanda L Tan, Karthik Gangavarapu, Michael R Wiley, Stephen White, Julien Thézé, Diogo M Magnani, Karla Prieto, Daniel Reyes, Andrea Bingham, Lauren M Paul, Refugio Robles-Sikisaka, Glenn Oliveira, Darryl Pronty, Hayden C Metsky, Mary Lynn Baniecki, Kayla G Barnes, Bridget Chak, Catherine A Freije, Adrianne Gladden-Young, Andreas Gnirke, Cynthia Luo, Bronwyn MacInnis, Christian B Matranga, Daniel J Park, James Qu, Stephen F Schaffner, Christopher Tomkins-Tinch, Kendra L West, Sarah M Winnicki, Shirlee Wohl, Nathan L Yozwiak, Joshua Quick, Joseph R Fauver, Kamran Khan, Shannon E Brent, Robert C Reiner, Paola N Lichtenberger, Michael Ricciardi, Varian K Bailey, David I Watkins, Marshall R Cone, Edgar W Kopp, Kelly N Hogan, Andrew C Cannons, Reynald Jean, Robert F Garry, Nicholas J Loman, Nuno R Faria, Mario C Porcelli, Chalmers Vasquez, Elyse R Nagle, Derek AT Cummings, Danielle Stanek, Andrew Rambaut, Mariano Sanchez-Lockhart, Pardis C Sabeti, Leah D Gillis, Scott F Michael, Trevor Bedford, Oliver G Pybus, Sharon Isern, Gustavo Palacios, Kristian G Andersen

**Affiliations:** Department of Immunology and Microbial Science, The Scripps Research Institute, La Jolla, CA 92037, USA; Center for Genome Sciences,U.S. Army Medical Research Institute of Infectious Diseases, Fort Detrick, MD 21702, USA; Department of Zoology, University of Oxford, Oxford OX1 3PS, UK; 4Vaccine and Infectious Disease Division, Fred Hutchinson Cancer Research Center, Seattle, WA 98109, USA; Department of Biological Sciences, College of Arts and Sciences, Florida Gulf Coast University, Fort Myers, FL, 33965,USA; Bureau of Public Health Laboratories, Division of Disease Control and Health Protection, Florida Department of Health, Miami, FL, 33125,USA; Department of Pathology, University of Miami Miller School of Medicine, Miami, FL, 33136, USA; Bureau of Epidemiology, Division of Disease Control and Health Protection, Florida Department of Health, Tallahassee, FL, 32399, USA; Scripps Translational Science Institute, La Jolla, CA 92037, USA; The Broad Institute of MIT and Harvard, Cambridge, MA 02142,USA; Institute of Microbiology and Infection, University of Birmingham, Birmingham B15 2TT, UK; Department of Microbiology, Immunology, and Pathology, Colorado State University, Fort Collins, CO 80523, USA; Li Ka Shing Knowledge Institute, St Michael´s Hospital, Toronto, ON M5B 1T8, Canada; Division of Infectious Diseases, Department of Medicine, University of Toronto, Toronto, ON M5B 1T8, Canada; Institute for Health Metrics and Evaluation, University of Washington, Seattle, WA 98121, USA; Division of Infectious Diseases, University of Miami Miller School of Medicine, Miami, FL, 33155, USA; Bureau of Public Health Laboratories, Division of Disease Control and Health Protection, Florida Department of Health, Tampa, FL, 33612, USA; Florida Department of Health in Miami-Dade County, Miami, FL, 33125, USA; Department of Microbiology and Immunology, Tulane University School of Medicine, New Orleans, LA 70112, USA; Miami-Dade County Mosquito Control, Miami, FL, 33178 USA; Department of Biology and Emerging Pathogens Institute, University of Florida, Gainesville, FL, 32610, USA; Institute of Evolutionary Biology, University of Edinburgh, Edinburgh EH9 3FL, UK; Fogarty International Center, National Institutes of Health, Bethesda, MD 20892, USA; Department of Integrative Structural and Computational Biology, The Scripps Research Institute, La Jolla, CA 92037, USA

## Abstract

Zika virus (ZIKV) is causing an unprecedented epidemic linked to severe congenital syndromes^1,2^. In July 2016, mosquito-borne ZIKV transmission was first reported in the continental United States and since then, hundreds of locally-acquired infections have been reported in Florida^3^. To gain insights into the timing, source, and likely route(s) of introduction of ZIKV into the continental United States, we tracked the virus from its first detection in Miami, Florida by direct sequencing of ZIKV genomes from infected patients and *Aedes aegypti* mosquitoes. We show that at least four distinct ZIKV introductions contributed to the outbreak in Florida and that local transmission likely started in the spring of 2016 - several months before its initial detection. By analyzing surveillance and genetic data, we discovered that ZIKV moved among transmission zones in Miami. Our analyses show that most introductions are phylogenetically linked to the Caribbean, a finding corroborated by the high incidence rates and traffic volumes from the region into the Miami area. By comparing mosquito abundance and travel flows, we describe the areas of southern Florida that are especially vulnerable to ZIKV introductions. Our study provides a deeper understanding of how ZIKV initiates and sustains transmission in new regions.

---

ZIKV transmission in the Americas was first reported in Brazil in May 2015^4^, though the virus was likely introduced 1-2 years prior to its detection^5^. By January 2016, the ZIKV epidemic had spread to several South and Central American countries and most islands in the Caribbean^6^. Like dengue virus (DENV) and chikungunya virus (CHIKV), ZIKV is vectored primarily by *Aedes* mosquitoes^7-10^. The establishment of the peridomestic species *Ae. aegypti* in the Americas^11^ has facilitated DENV, CHIKV, and now likely ZIKV to become endemic in this region^12^. In the continental United States, transient outbreaks of DENV and CHIKV have been reported in regions of Texas and Florida^13-19^ with abundant seasonal *Ae. aegypti* populations^11,20^.

In July 2016, the first locally-acquired ZIKV cases in the continental United States were reported in Florida^3^. The outbreak, which generated 256 confirmed ZIKV infections in 2016, was likely initiated by travel-associated infections in Florida (Fig. 1a). While transmission was confirmed across four counties in Florida (Fig. 1b), the ZIKV outbreak was most intense in Miami-Dade County (241 infections). Although the location of exposure could not be determined in all cases, at least 114 (47%) of the infections were likely acquired in one of three distinct transmission zones within Miami: Wynwood, Miami Beach, and Little River (Fig. 1c-d).

**Figure 1.**
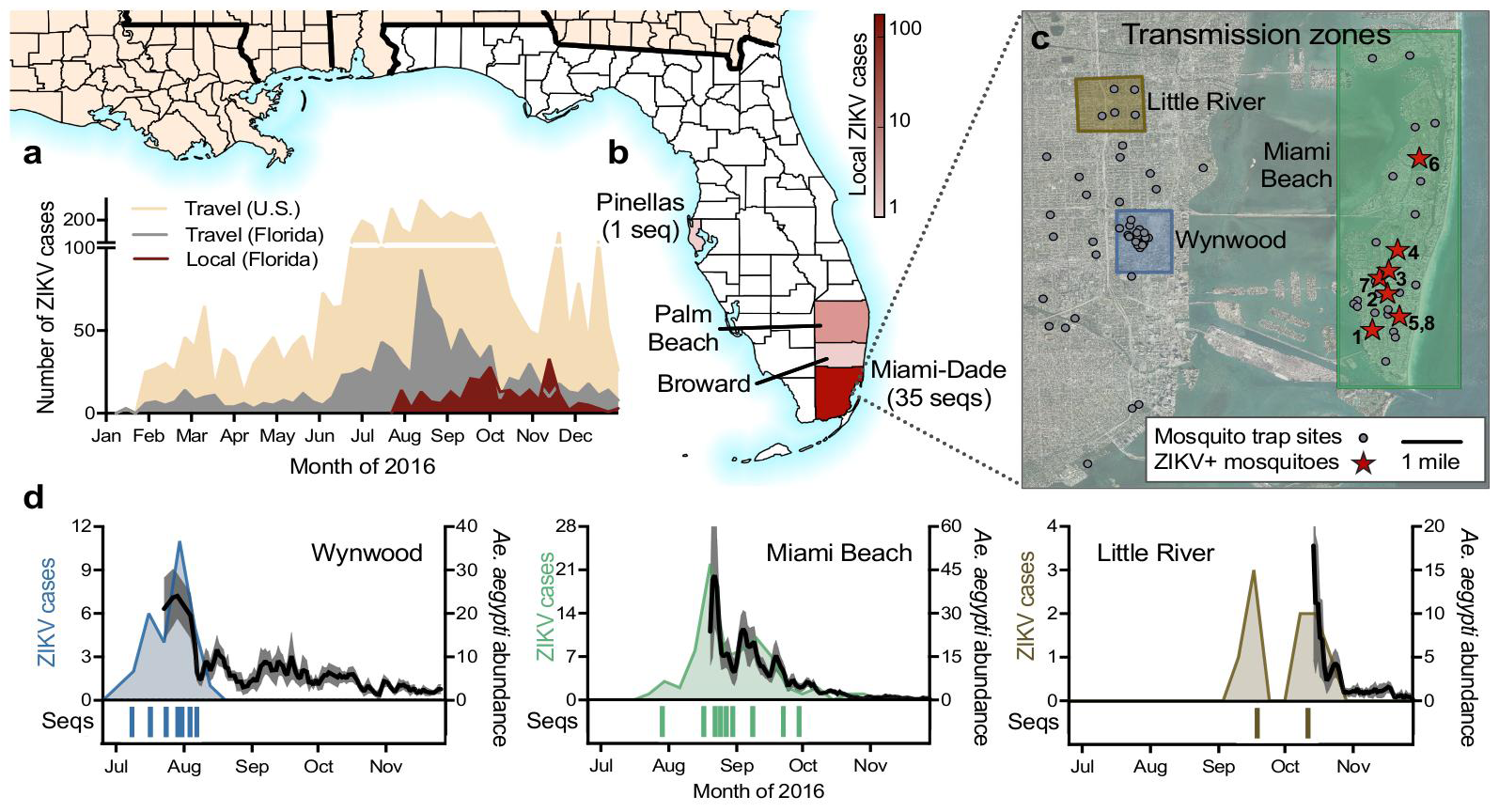
Zika virus outbreak in Florida. **(a)** Weekly counts of confirmed travel-associated and locally-acquired ZIKV cases in 2016. **(b)** Four counties reported locally-acquired ZIKV cases in 2016: Miami-Dade (241), Broward (5), Palm Beach (8), Pinellas (1), and unknown origin (1). **(c)** The locations of mosquito traps and collected *Ae. aegypti* mosquitoes found to contain ZIKV RNA (ZIKV+) in relation to the transmission zones within Miami. **(d)** Temporal distribution of weekly ZIKV cases (left y-axis), sequenced cases (bottom), and *Ae. aegypti* abundance per trap night (right y-axis) associated with the three described transmission zones. ZIKV cases and sequences are plotted in relation to symptom onset dates (n=18). Sequenced cases without onset dates or that occurred outside of the transmission zones are not shown (n=10). Human cases and *Ae.aegypt* i abundance per week werepositively correlated (Spearman r = 0.61, Supplementary Data).

Using mosquito surveillance data, we determined the extent of mosquito-borne ZIKV transmission in Miami. Of the 24,351 mosquitoes collected throughout Miami-Dade County from June to November 2016, 99.8% were *Ae. aegypti* and 8 pools of *Ae. aegypti* collected in Miami Beach tested positive for ZIKV RNA (Fig. 1c, Supplementary Fig. 1). From these pools, we estimated that ∼1 out of 1,600 *Ae. aegypti* mosquitoes were infected (0.061%, 95% CI: 0.028-0.115%, Supplementary Fig. 1a). This is similar to reported *Ae. aegypti* infection rates during DENV and CHIKV outbreaks^21^. Even though ZIKV-infected mosquitoes were not detected outside Miami Beach (Fig. 1c), we found that the number of human ZIKV cases correlated strongly with *Ae. aegypti* abundance within each of the defined transmission zones (Spearman r = 0.61, Fig. 1d, Supplementary Fig. 1b). This suggests that *Ae. aegypti* mosquitoes were the primary mode of transmission throughout Miami-Dade County and that changes to vector abundance directly impacted human infection rates. The application of insecticides^3^, which we found suppressed local mosquito populations during the periods of intensive usage (Supplementary Fig. 1c), therefore likely contributed to the clearance of ZIKV within these zones.

To gain molecular insights into the Florida outbreak, we sequenced 39 ZIKV genomes directly from clinical and mosquito samples without cell culture enrichment^22^ (Supplementary File 1). Our Florida ZIKV dataset (36 genomes) included 29 genomes from patients with locally-acquired infections (Fig. 1d) and 7 from *Ae. aegypti* mosquito pools (Fig. 1c). We also sequenced 3 ZIKV genomes from travel-associated cases diagnosed in Florida. Our dataset included cases from all three transmission zones in Miami (Fig. 1d) and represented ∼11% of all confirmed locally-acquired casesin Florida. We made all sequence data openly available at NCBI (BioProjects PRJNA342539, PRJNA356429) and other online resources immediately after data generation.

To infer viral genealogies, we reconstructed phylogenetic trees from our 39 ZIKV genomes along with 65 published genomes from other affected regions (Fig. 2, Supplementary Fig. 2 and 3). We found that the Florida ZIKV genomes formed four distinct lineages (labeled F1-F4, Fig. 2a), three of which (F1-F3) belonged to the same major clade (labeled A, Fig. 2a). We only sampled a single human case each from the F3 and F4 lineages, consistent with limited transmission (Fig. 2a). The other two Florida lineages (F1-F2) comprised ZIKV genomes sequenced from multiple human and mosquito samples within Miami-Dade County (Fig. 2b).

**Figure 2.**
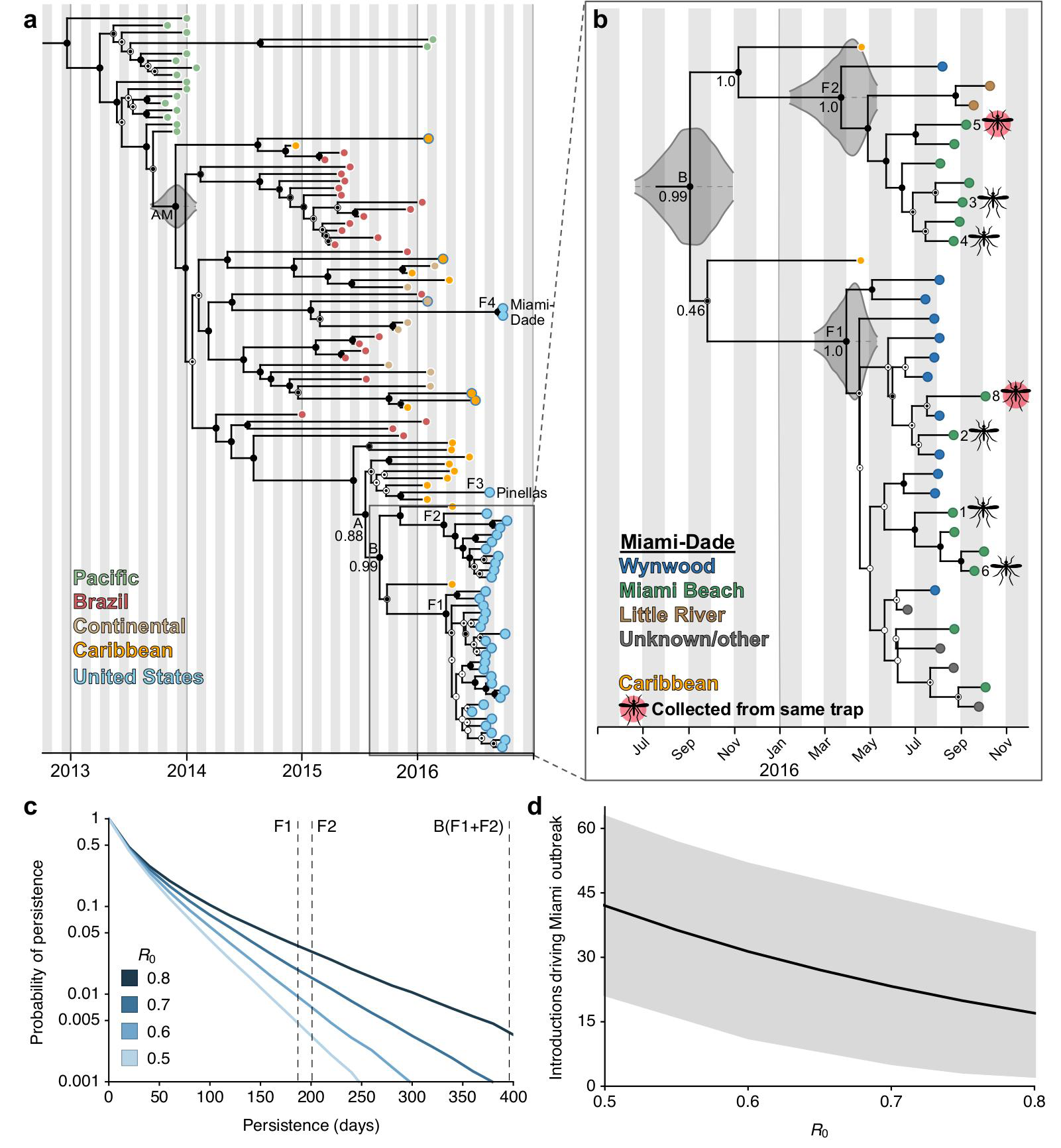
Multiple introductions of Zika virus into Florida. (**a**) Maximum clade credibility (MCC) tree of ZIKV genomes sequenced from outbreaks in the Pacific islands and the epidemic in the Americas. Tips are colored based on collection location. The five tips outlined in blue but filled with a different color indicate ZIKV cases in the United States associated with travel (fill color indicates the probable location of infection). Clade posterior probabilities are indicated with black circles inside white circles with posterior probability of 1 filling the entire circle black. The grey violin plot indicates the 95% highest posterior density (HPD) interval for the tMRCA for the epidemic in the Americas (AM)^5^. Lineage F4 contains two identical ZIKV genomes from the same patient. (**b**) A zoomed in version of the whole MCC tree showing the collection locations of Miami-Dade sequences and whether they were sequenced from mosquitoes (numbers correspond to trap locations in Fig. 1c). 95% HPD intervals are shown for the tMRCAs (**c**) The probability of ZIKV persistence after introduction for different *R*_0_. Persistence is measured as the number of days from initial introduction of viral lineages until their extinction. Vertical dashed lines show the inferred mean persistence time for lineages F1, F2 and B based on their tMRCA. (**d**) Total number of introductions (mean with 95% CI) that contributed to the outbreak of 241 local cases in Miami-Dade County for different *R*_0_.

Using time-structured phylogenies^23^, we estimated that at least four separate introductions were responsible for the locally-acquired cases observed in our dataset. The phylogenetic placement of lineage F4 clearly indicates that it resulted from an independent introduction of a lineage genetically distinct from those in clade A (Fig. 2a). For the two well-supported nodes linking lineages F1 and F2 (labeled B, Fig. 2a) and F1-F3 (A, Fig. 2a), we estimated the time of the most recent common ancestor (tMRCA) to be during the summer of 2015. Our data displayed a strong clock signal (Supplementary Fig. 2b) and tMRCA estimates were robust across a range of different molecular clock and coalescent models (Supplementary Table 1). Thus while F1-F3 all belong to clade A, any less than three distinct introductions leading to these lineages would have required undetected transmission of ZIKV in Florida for approximately one year (Fig. 2a).

To estimate the likelihood of a single ZIKV transmission chain persisting for over a year, we modeled epidemic spread under different assumptions of the basic reproductive number (*R*_0_). We calculated *R*_0_ using the locally-acquired and travel-associated case counts in Miami-Dade County, along with the number of observed genetic lineages. Allowing for variability in the percentage of infectious travel-associated cases and for sampling bias across lineages, we estimated an *R*_0_ for Miami-Dade between 0.5 and 0.8 (Supplementary Fig. 4). Even at the upper end of this range, the probability of a single transmission chain persisting for over a year is extremely low (∼0.5%, Fig. 2c). This is especially true considering the low *Ae. aegypti* abundance during the winter months (Supplementary Fig. 1d). Given the low probability of long-term persistence, we expect that the ZIKV genomes we sequenced from Florida (F1-F4) were the result of at least four separate introductions. Differences in surveillance practices across areas, combined with the high number of travel-associated cases in Florida (Fig. 1a), however, likely mean that unsampled ZIKV introductions contributed to the outbreak. To estimate the total number of underlying ZIKV introductions, we modeled epidemic scenarios that resulted in 241 locally-acquired cases within Miami-Dade County, and found that with *R*_0_ values of 0.5-0.8, we expect 17-42 (95% CI 3-63) separate introductions to have contributed to the Miami-Dade outbreak (Fig. 2d). The majority of these introductions would likely have generated a single secondary case that was undetected in our genetic sampling (Supplementary Fig. 4a).

The two main ZIKV lineages, F1 and F2, included the vast majority of genomes we sampled in Florida (92%, Fig. 2a). Assuming they represent two independent introductions, we estimated when each of these lineages arrived in Florida. The probability densities for the tMRCAs of both F1 and F2 were centered around March-April, 2016 (Fig. 2b, 95% highest posterior density [HPD]: January-May, 2016). The estimated timing for these introductions corresponds with suitable *Ae. aegypti* populations in Miami-Dade County^24^ (Supplementary Fig. 1d) and suggests that ZIKV transmission in Florida could have started at least two months prior to its first detection in July 2016 (Fig. 1a). The dates of the introductions could be more recent if multiple F1 or F2 lineage viruses arrived into Florida independently. However, more than 2 introductions from each lineage would be necessary to substantially change our estimates for the timing of the earliest introduction of ZIKV into Florida.

To better understand the transmission dynamics within Miami, we analyzed our genomic data together with case investigation data from the Florida Department of Health (DOH, Supplementary Table 1). While spatially distinct, the three ZIKV transmission zones occurred within ∼3 miles of each other (Fig. 1c) and we found that the ZIKV infections associated with each zone overlapped temporally (Fig. 1d). Our 22 ZIKV genomes with zone assignments all belonged to lineages F1 and F2, but neither of these lineages were confined to a single transmission zone (Fig. 2b). In fact, we detected both F1 and F2 lineage viruses from *Ae. aegypti* collected from the same trap 26 days apart (mosquitoes 5 and 8, Fig. 2b). The detection of genetically similar viruses and the temporal overlap of ZIKV infections suggest that ZIKV moved among areas of Miami.

Given that ZIKV transmission is unlikely to persist through the winter in Florida^25^ (Fig. 2c, Supplementary Fig. 1d), determining the sources and routes of ZIKV introductions could help mitigate future outbreaks. We found that lineages F1-F3 clustered with ZIKV genomes sequenced from the Dominican Republic and Guadeloupe (Fig. 2, Supplementary Fig. 2 and 3). In contrast, the F4 lineage clustered with genomes from Central America (Fig. 2, Supplementary Fig. 2 and 3). These findings suggest that while ZIKV outbreaks occurred throughout the Americas, the Caribbean islands appear to have been the main source of ZIKV that resulted in local transmission in Florida. Because of the severe undersampling of ZIKV genomes from across the Americas, however, we cannot rule out other source areas or panmixis of the ZIKV population in the region. Similarly, even though we found that the Florida ZIKV genomes clustered together with sequences from the Dominican Republic, our results do not prove that ZIKV entered Florida from this country.

We investigated ZIKV infection rates and travel patterns to corroborate our phylogenetic evidence for Caribbean introductions into Florida. We found that the Caribbean islands bore the highest ZIKV incidence rates around the likely time of the main introductions into Florida (Fig. 2b), despite Brazil and Colombia reporting the highest absolute number of ZIKV cases (January to June, 2016, Fig. 3a, Supplementary Fig. 5, Supplementary File 2). During the same time period, ∼3 million travelers arriving from the Caribbean accounted for 54% of the total traffic into Miami, with the vast majority (∼2.4 million) arriving via cruise ships (Fig. 3b, Supplementary Fig. 6, Supplementary File 2). Combining the infection rates with travel capacities, we estimated that ∼60-70% of ZIKV infected travelers entering Miami arrived from the Caribbean (Fig. 3c and Supplementary Fig. 7a). In support of these estimates, we found that the observed number of travel-associated ZIKV cases correlated strongly with the expected number of importations from the Caribbean (Spearman r = 0.8, Fig. 3d, Supplementary Fig. 7b). Also, 67% of the travel-associated infections diagnosed in Florida reported recent travel to the Caribbean (Fig. 3e); however, the mode of travel for these cases is unknown. Taken together, these findings suggest that a high incidence of ZIKV in the Caribbean, combined with frequent travel from the region, could have played a key role in the establishment of ZIKV transmission in Florida in the spring of 2016. These findings, however, do not indicate that cruise ships themselves are risk factors for human ZIKV infection, but only that they served as a major mode of transportation from areas with active transmission. In addition, the probability of ZIKV exposure may vary among individuals depending on their purpose of travel and therefore, we cannot determine the specific contribution of ZIKV-infected travelers arriving via airlines or cruise ships.

**Figure 3.**
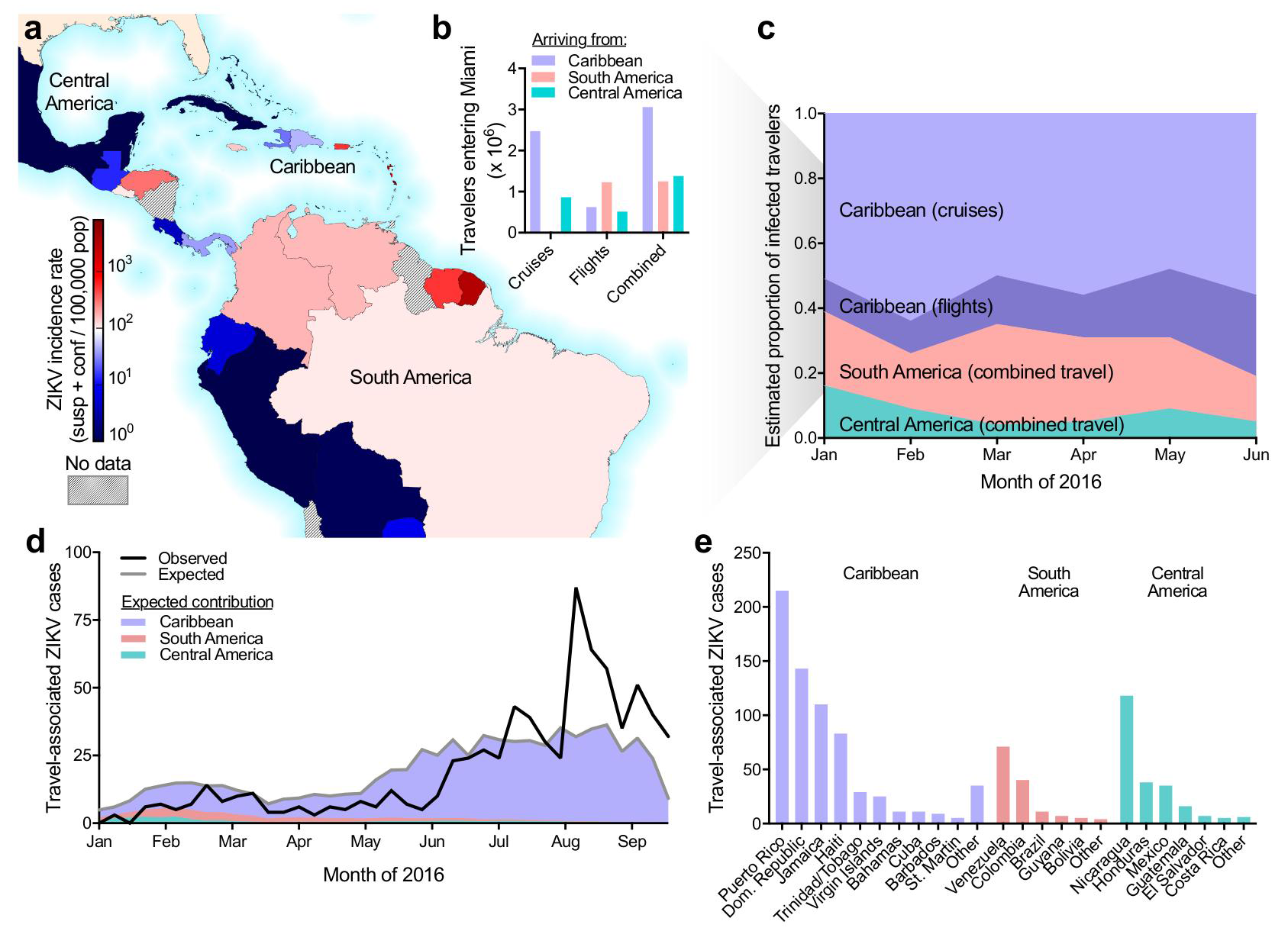
Frequent opportunities for Zika virus introductions into Miami from the Caribbean. (**a**) Reported ZIKV cases per country/territory from January to June, 2016 normalized by total population. (**b**) The number of estimated travelers entering Miami during January to June, 2016 by method of travel. (**c**) The number of travelers and the reported ZIKV incidence rate for the country/territory of origin were used to estimate the proportion of infected travelers coming from each region with ZIKV in the Americas. (**d**) The observed number of weekly travel-associated ZIKV cases in Florida were plotted with the expected number of ZIKV-infected travelers (as estimated in panel c) coming from all of the Americas (grey line) and the regional contributions (colored areas). (**e**) The countries visited by the 1,016 travel-associated ZIKV cases diagnosed in Florida.

The vast majority of the Florida ZIKV outbreak occurred in Miami-Dade County (94% of reported local cases in Florida, Fig. 1b). To determine if there is a higher potential for ZIKV outbreaks in the Miami area, we analyzed incoming passenger traffic by air and sea from regions with ZIKV transmission and *Ae. aegypti* abundance. We found that Miami and nearby Fort Lauderdale received ∼72% of the traffic entering Florida (Fig. 4) and Miami received more air and sea traffic than any other city in the continental United States in 2016 (Supplementary Fig. 8). During January to April 2016, we estimated that *Ae. aegypti* abundance was highest in southern Florida^20^ (Fig. 4, Supplementary Fig. 1d, Supplementary Fig. 8). By June, most of Florida and several cities across the South, such as New Orleans, Atlanta, and Charleston, were expected to support high seasonal *Ae. aegypti* populations^20^ (Supplementary Fig. 8); however, none of these cities have reported local *Ae. aegypti*-borne transmission of any virus in at least 60 years^17^. In fact, despite the thousands of opportunities for introductions across the country (Fig. 1a), the only region outside of Florida that has reported mosquito-borne ZIKV transmission is southern Texas^27^. Southern Texas is also the only other region to recently report DENV outbreaks^17-19^. Therefore, the combination of international travelers, mosquito ecology, and human population density near Miami likely make it one of the few places in the continental United States most at risk for *Ae. aegypti*-borne virus outbreaks^20,24,26^.

**Figure 4.**
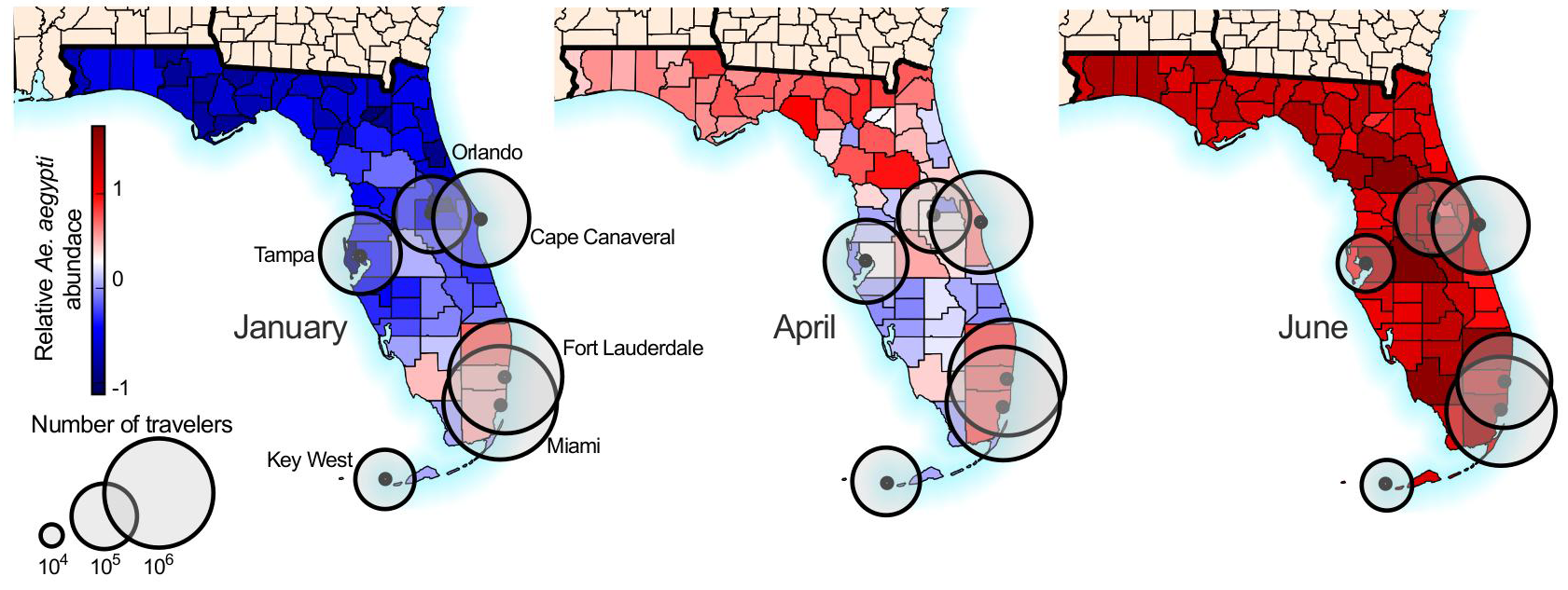
Southern Florida has a high potential for Aedes aegypti-borne virus outbreaks. The number of travelers per month (circles) entering Florida cities via flights and cruise ships were plotted with estimated relative *Ae. aegypti* abundance. Only cities receiving >10,000 passengers per month are shown. Relative *Ae. aegypti* abundance for every month is shown in Supplementary Fig. 1d.

The extent of ZIKV transmission in Florida was unprecedented, with more reported ZIKV cases in 2016 (256) than DENV cases since 2009 (136)^13^. Given that the majority of ZIKV infections are asymptomatic^2,28^, the true number of ZIKV cases was likely much higher. Despite this, we estimated that the average *R*_0_ was less than 1 and therefore multiple introductions were necessary to give rise to the magnitude of the observed outbreak^25^. The high volume of airline and cruise ship traffic entering Florida from ZIKV-affected regions, especially the Caribbean, likely provided a substantial supply of ZIKV-infected individuals^29,30^. Because Florida is unlikely to sustain long-term ZIKV transmission^25^, the potential for future ZIKV outbreaks in this region is highly dependent upon activity elsewhere. Therefore, we expect that outbreaks in Florida will cycle with the ZIKV transmission dynamics in the Americas^12^.

## Acknowledgements

The authors thank Chet Moore, Barry Alto, and Sophie Taylor for discussions, Andrew Monaghan for providing details about the *Ae. aegypti* abundance data from United States port cities, Marshall Pilcher from ThermoFisher for sequencing assistance, and Gary P. Schroth and Stephen M. Gross from Illumina for designing and providing the probes used for enrichment-based sequencing. N.D.G. is supported by a National Institutes of Health (NIH) training grant 5T32AI007244-33. G.D. is supported by the Mahan Postdoctoral Fellowship from the Computational Biology Program at Fred Hutchinson Cancer Research Center. D.A.T.C. was supported by the US NIH MIDAS program (U54-GM088491) as well as Cooperative Agreement U01CK000510 funded by the Centers for Disease Control and Prevention (CDC). A.R. is supported by the European Union Seventh Framework Programme (FP7/2007-2013) under Grant Agreement 278433-PREDEMICS and ERC Grant agreement 260864 and Horizon 2020 research and innovation program Grant Agreement 643476-COMPARE. T.B. is a Pew Biomedical Scholar and his work is supported by NIH award R35 GM119774-01. O.G.P. received funding from the European Research Council under the European Union’s Seventh Framework Programme (FP7/2007-2013)/ERC, grant agreement number 614725-PATHPHYLODYN and generous support of the American people through the United States Agency for International Development Emerging Pandemic Threats Program-2 PREDICT-2 (Cooperative Agreement No. AID-OAA-A-14-00102). S.I. and S.F.M. are supported by a NIH National Institute of Allergy and Infectious Diseases (NIAID) award 4R01AI099210-04. ZIKV sequencing at the US Army Medical Research Institute of Infectious Diseases (USAMRIID) was supported by the Defense Advanced Research Projects Agency (DARPA). K.G.A. is a Pew Biomedical Scholar, and his work is supported by an NIH National Center for Advancing Translational Studies Clinical and Translational Science Award UL1TR001114 and NIAID contract HHSN272201400048C. The content of this publication does not necessarily reflect the views or policies of the US Army, the Department of Health and Human Services, the CDC, or the Florida DOH.

## Author Contributions

All contributions are listed in order of authorship. Designed the experiments: N. D. G., J. T. L., G. D., M. U. G. K., D. A. T. C., P. C. S, L. D. G., S. F. M., T. B., O. G. P., S. I., G. P., and K. G. A.; Collected samples: A. L. T., S. W., D. M. M., A. B., L. M. P., D. P., P. N. L., M. R., V. K. B., D. I. W., M. R. C., E. W. K., K. N. H., A. C. C., R. J., M. C. P., C. V., D. S., L. D. G., S. F. M., and S. I.; Performed the sequencing: N. D. G., M. W. R., K. P., D. R., R. R.-S., G. O., and E. N.; Provided data, critical reagents, or protocols: N. D. G., J. T. L., G. D., M. U. G. K., K. G., M. R. W., R. R.-S., G. O., H. C. M., M. L. B., K. G. B., B. C., C. A. F., A. G.-Y., A. G., C. L., B. M., C. B. M., D. J. P., J. Q., S. F. S., C. T.-T., K. L. M., S. M. W., S. W., N. L. Y., J. Q., J. R. F., K. K., S. E. B., R. F. G., N. J. L., M. C. P., C. V., P. C. S., S. F. M., and S. I.; Analyzed the data: N. D. G., J. T. L., G. D., M. U. G. K., K. G., J. T., J. R. F., R. C. R., N. R. F., D. A. T. C., A. K., M. S.-L., T. B., S. F. M, O. G. P., S. I., and K. G. A.; Edited the manuscript: G. D., M. U. G. K., J. T., S. F. S., A. R., T. B., O. G. P., S. I., and G. P.; Wrote the manuscript: N. D. G., J. T. L., and K. G. A.; All authors read and approved the content of the manuscript.

## Methods

### Ethical statement

This study has been evaluated and determined to be exempt by the Institutional Review Boards (IRB) at The Scripps Research Institute (TSRI) and the USAMRIID Office of Human Use and Ethics. The study was reviewed by the Florida DOH Human Research Protection Program and determined not to require IRB review.

### Florida Zika virus case data

Weekly reports of international travel-associated and locally-acquired ZIKV infections diagnosed in Florida were obtained from the Florida DOH mosquito-borne disease surveillance system^13^. Dates of symptom onset from the Miami transmission zones (Wynwood, Miami Beach, and Little River) determined by the Florida DOH investigation process were obtained from the ZIKV resource website^31^ and daily updates^32^. International travel-associated ZIKV case counts in the United States (outside of Florida) were obtained from the CDC^33^. The local and travel-associated ZIKV case numbers for Florida were obtained from the Florida DOH. The one local ZIKV infection diagnosed in Duval County was believed to have originated elsewhere in Florida. Therefore, this case is listed as “unknown origin” in Fig. 1b. In Fig. 3e, only the countries visited 5 or more times by ZIKV-infected travelers diagnosed in Florida are shown. Countries with 5 or less visits were aggregated into an “other” category by region (*i.e.*, Caribbean, South America, or Central America).

### Clinical sample collection

Clinical samples from locally-acquired ZIKV infections were collected from June 22 to October 11, 2016. The Florida DOH identified persons with compatible illness and clinical samples were shipped to the Bureau of Public Health Laboratories for confirmation by qRT-PCR and antibody tests following interim guidelines^3,34-36^. Clinical specimens (whole blood, serum, saliva, or urine) submitted for analysis were refrigerated or frozen at ≤ −70°C until RNA was extracted. RNA was extracted using the RNAeasy kit (QIAGEN), MagMAX for Microarrays Total RNA Isolation Kit (Ambion), or MagNA Pure LC 2.0 or 96 Systems (Roche Diagnostics). Purified RNA was eluted into 50-100 µL using the supplied elution buffers, immediately frozen at ≤ −70°C, and transported on dry ice. The Florida DOH also provided investigation data for these samples, including symptom onset dates and, when available, assignments to the zone where infection likely occurred (Supplementary File 1).

### Mosquito collection and entomological data analysis

24,351 *Ae. aegypti* and *Ae. albopictus* mosquitoes (sorted into 2,596 pools) were collected throughout Miami-Dade County during June to November, 2016 using BG-Sentinel mosquito traps (Biogents AG). Up to 50 mosquitoes of the same species and sex were pooled per trap. The pooled mosquitoes were stored in RNAlater (Invitrogen), RNA was extracted using either the RNAeasy kit (QIAGEN) or MagMAX for Microarrays Total RNA Isolation Kit (Ambion), and ZIKV RNA was detected by qRT-PCR targeting the envelope protein coding region^36^ or the Trioplex qRT-PCR kit^37^. ZIKV infection rates were calculated per 1,000 female *Ae. aegypti* mosquitoes using the bias-corrected maximum likelihood estimate (MLE)^38^. Days of insecticide usage by the Miami-Dade Mosquito Control were inferred from the zone-specific ZIKV activities timelines published by the Florida DOH^31^.

### Relative monthly Ae. aegypti abundance

For the purpose of this study we used *Ae. aegypti* suitability maps from Kraemer *et al*.^11^ and derived monthly estimates based on the statistical relationships between mosquito presence and environmental correlates^41^. Following Hwang *et al*.^42^ we used a simple mathematical formula to transform the probability of detection maps into mosquito abundance maps. In order to do so, we assumed P (Y=1) where Y is a binary variable (presence/absence). Using a Poisson distribution X() to govern the abundance of mosquitoes, the probability of not observing any mosquitoes can be related to the probability of absence as: P(X=0)=P(Y=0). We used the following transformation to generate abundance (λ) estimates per county in Florida:

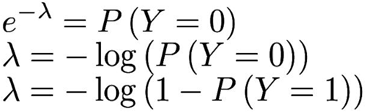

### Zika virus quantification

ZIKV genome equivalents (GE) were quantified by qRT-PCR. At TSRI, ZIKV qRT-PCR was performed as follows: ZIKV RNA standards were transcribed from the ZIKV NS5 region (8651-9498 nt) using the T7 forward primer (5’ - TAA TAC GAC TCA CTA TAG GGA GA TCA GGC TCC TGT CAA AAC CC - 3’), reverse primer (5’ - AGT GAC AAC TTG TCC GCT CC - 3’), and the T7 Megascript kit (Ambion). For qRT-PCR, primers and a probe targeting the NS5 region (9014-9123 nt) were designed using the ZIKV isolate PRVABC59 (GenBank: KU501215): forward primer (5’- AGT GCC AGA GCT GTG TGT AC - 3’), reverse primer (5’ - TCT AGC CCC TAG CCA CAT GT - 3’), and FAM-fluorescent probe (5’ - GGC AGC CGC GCC ATC TGG T - 3’). We have the best probes. The qRT-PCR assays were performed in 25 µl reactions using the iScript One-step RT-PCR Kit for probes (Bio-Rad Laboratories Inc.) and 2 µl of sample RNA. Amplification was performed at 50°C for 20 min, 95°C for 3 min, and 40 cycles of 95°C for 10 s and 57°C for 10 s. Fluorescence was read at the end of the 57°C annealing-extension step. 10-fold dilutions of the ZIKV RNA transcripts (2 µl/reaction) were used to create a standard curve for quantification of ZIKV GE/µl of RNA.

ZIKV GE were quantified at USAMRIID using the University of Bonn ZIKV envelope protein (Bonn E) qRT-PCR assay^43^. RNA standards were transcribed using an amplicon generated from a ZIKV plasmid containing T7 promoter at the start of the 5’ untranslated region (UTR). The plasmid was designed using the ZIKV isolate BeH819015 (GenBank: KU365778.1) and the amplicon included nts 1-4348, which covers the 5’ UTR, C, prM, M, E, NS1, and NS2 regions. The qRT-PCR assays were performed in 25 µl reactions using the SuperScript III platinum One-step qRT-PCR Kit (ThermoFisher) and 2 µl of sample RNA was used. Amplification was performed following conditions as previously described^43^. 10-fold dilutions of the ZIKV RNA transcripts (5 µl/reaction) were used to create a standard curve for quantification of ZIKV GE/µl of RNA.

### Amplicon-based Zika virus sequencing

ZIKV sequencing at TSRI was performed using an amplicon-based approach using the ZikaAsian V1 scheme, as described^22^. Briefly, cDNA was reverse transcribed from 5 µl of RNA using SuperScript IV (Invitrogen). ZIKV cDNA (2.5 µl/reaction) was amplified in 35 × 400 bp fragments from two multiplexed PCR reactions using Q5 DNA High-fidelity Polymerase (New England Biolabs). The amplified ZIKV cDNA fragments (50 ng) were prepared for sequencing using the Kapa Hyper prep kit (Kapa Biosystems) and SureSelect XT2 indexes (Agilent). Agencourt AMPure XP beads (Beckman Coulter) were used for all purification steps. Paired-end 251 nt reads were generated on the MiSeq using the V2 500 cycle or V3 600 cycle kits (Illumina).

Trimmomatic was used to remove primer sequences (first 22 nt from the 5’ end of the reads, which is the maximum length of the primers used for the multiplexed PCR) and bases at both ends with Phred quality score < 20^44^. The reads were then aligned to the complete genome of a ZIKV isolate from the Dominican Republic, 2016 (GenBank: KU853012) using Novoalign v3.04.04 (www.novocraft.com). Samtools was used to sort the aligned BAM files and to generate alignment statistics^45^. Snakemake was used as the workflow management system^46^. The code and reference indexes for the pipeline can be found at https://github.com/andersen-lab/zika-pipeline. ZIKV-aligned reads were visually inspected using Geneious v9.1.5^47^ before generating consensus sequences. A minimum of 3× read-depth coverage, in support of the consensus, was required to make a base call.

The consensus ZIKV sequences from FL01M and FL03M generated by sequencing 35 × 400 bp amplicons on the MiSeq were validated using the following approaches: 1) sequencing the 35 × 400 bp amplicons on the Ion S5 platform (ThermoFisher), 2) sequencing amplicons generated using an Ion AmpliSeq^®^ (ThermoFisher) panel customly targeted towards ZIKV on the Ion S5 platform, and 3) sequencing 5 × 2,150-2,400 bp ZIKV amplicons on the MiSeq. For Ion library preparation, cDNA was synthesized using the SuperScript VILO kit (ThermoFisher). ThermoFisher designed 875 custom ZIKV primers to produce 75 amplicons of ∼200 bp in two PCR reactions for use with their Ion AmpliSeq Library Kit 2.0. The reagent FuPa was used to digest the modified primer sequences after amplification. The DNA templates were loaded onto Ion 520 chips using the Ion Chef and sequenced on the Ion S5 with the 200 bp output (ThermoFisher). The 35 × 400 bp amplicons generated for the MiSeq as described above were introduced into the Ion workflow using the Ion AmpliSeq Library Kit 2.0, but without fragmentation. Primers to amplify 2,150-2,400 bp ZIKV fragments (Supplementary File 3) were kindly provided by Shelby O’Connor, Dawn Dudly, Dave O’Connor, and Dane Gellerup (AIDS Vaccine Research Laboratory, University of Wisconsin, Madison). Each fragment was amplified individually by PCR using the cDNA generated above, Q5 DNA High-fidelity Polymerase, and the following thermocycle conditions: 55 ℃ for 30 m, 94 ℃ for 2 m, 35 cycles of 94 ℃ for 15 s, 56 ℃ for 30 s, and 68 ℃ for 3.5 m, 68 ℃ for 10 m, and held at 4 ℃ until use. Each PCR product was purified using Agencourt AMPure XP beads, sheared to 300 to 400 nt fragments using the Covaris S2 sonicator, indexed and prepared for sequencing as described above, and sequenced using the MiSeq V2 500 cycle kit (paired-end 251 nt reads). Compared to the consensus sequences generated using 35 × 400 bp amplicons on the MiSeq, there were no consensus-level mismatches in the coding sequence using any of the other three approaches (Supplementary Table 2). There were, however, some mismatches in the 5’ and 3’ UTRs (where the genomic RNA is heavily structured), likely a result of PCR bias and decreased coverage depth.

### Enrichment-based Zika virus sequencing

ZIKV sequencing at USAMRIID was performed using a targeted enrichment approach. Sequencing libraries were prepared using the TruSeq RNA Access Library Prep kit (Illumina) with custom ZIKV probes. The set included 866 unique probes each of which was 80 nt in length (Supplementary File 3). The probes were designed to cover the entire ZIKV genome and to encompass the genetic diversity present on GenBank on January 14, 2016. In total, 26 ZIKV sequences were used during probe design (Supplementary File 3). Extracted RNA was fragmented at 94 ˚C for 0-60 s and each sample was enriched separately using a quarter of the reagents specified in the manufacturer’s protocol. Samples were barcoded, pooled and sequenced using the MiSeq Reagent kit v3 (Illumina) on an Illumina MiSeq with a minimum of 2 × 151 bp reads. Dual indexing, with no overlapping indices, was used.

The random hexamer associated with read one and the Illumina adaptors were removed from the sequencing reads using Cutadapt v1.9.dev1^48^, and low-quality reads/bases were filtered using Prinseq-lite v0.20.3^49^. Reads were aligned to a reference genome (GenBank: KX197192.1) using Bowtie2 v2.0.6^50^, duplicates were removed with Picard (http://broadinstitute.github.io/picard), and a new consensus was generated using a combination of Samtools v0.1.18^45^ and custom scripts (https://github.com/jtladner/Scripts/blob/master/reference-based_assembly/consensus_fasta.py). Only bases with Phred quality score ≥ 20 were utilized in consensus calling, and a minimum of 3× read depth coverage, in support of the consensus, was required to make a call; positions lacking this depth of coverage were treated as missing (*i.e.* called as ‘‘N’’).

### Phylogenetic analyses

All published and available complete ZIKV genomes of the Asian genotype from the Pacific and the Americas were retrieved from GenBank public database as of December 2016. Public sequences (n=65) were codon-aligned together with ZIKV genomes generated in this study (n=39) using MAFFT^51^ and inspected manually. The multiple alignment contained 104 ZIKV 53 sequences collected between 2013 and 2016, from the Pacific (American Samoa, French Polynesia, and Tonga), Brazil, other South and Central Americas (Guatemala, Mexico, Suriname, and Venezuela), the Caribbean (Dominican Republic, Guadeloupe, Haiti, Martinique, and Puerto Rico), and the United States (Supplementary File 1).

In order to determine the temporal signal of the sequence dataset, a maximum likelihood (ML) phylogeny was first reconstructed with PhyML^52^ using the general time-reversible (GTR) nucleotide substitution model and gamma distributed rates amongst sites^54^ (Supplementary File 1), which was identified as the best fitting model for ML inference by jModelTest2^55^. Then, a correlation between root-to-tip genetic divergence and date of sampling was conducted in TempEst^56^.

Bayesian phylogenetic analyses were performed using BEAST v.1.8.4^23^ to infer time-structured phylogenies. We used an SDR06 nucleotide substitution model^57^ with a non-informative continuous time Markov chain reference prior (CTMC)^58^ on the molecular clock rate. Replicate analyses using multiple combinations of molecular clock and coalescent demographic models were explored to select the best fitting model by marginal likelihood comparison using path-sampling and stepping-stone estimation approaches^59-61^ (Supplementary Table 3). The best fit model was a strict molecular clock along with a Bayesian skyline demographic model^62^. All the Bayesian analyses were run for 30 million Markov chain Monte Carlo steps, sampling parameters and trees every 3000 generations (BEAST XML file and MCC tree available in Supplementary File 1).

### Expected number and distribution of local cases from Zika virus importations

We used branching process theory^63,64^ to generate the offspring distribution (subsequent local cases) that is expected from a single introduction. The offspring distribution *L* is modelled with a negative binomial distribution with mean *R*_0_ and over-dispersion parameter *k.* The total number of cases *j* that is caused by a single importation (including the index case) after an infinite time^65^ has the following form:

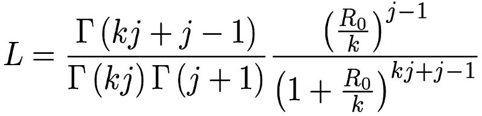

The parameter *k* represents the variation in the number of secondary cases generated by each case of ZIKV^63^. In the case of vector borne diseases, local heterogeneity is high due to a variety of factors such as mosquito population abundance, human to mosquito interaction, and control interventions^66,67^. Here, we assumed high heterogeneity (*k*=0.1) following previous estimates for vector borne diseases^64^. This distribution *L* is plotted in Supplementary Fig. 4a. For the following, we took a forward simulation approach, drawing random samples from this distribution. All estimates were based on 100,000 random simulations.

We used this formula to estimate the probability of observing 241 local cases in Miami-Dade County alongside 320 travel-associated cases. We approached this by sampling 320 introduction events from *L* and calculating the total number of local cases in the resulting outbreak (Supplementary Fig. 4b). We also calculated the likelihood of observing 241 local cases in the total outbreak (Supplementary Fig. 4c), finding that the MLE of *R*_0_ lies between 0.35 and 0.55. As a sensitivity analysis, we additionally modelled introductions with the assumption that only 50% of travelers were infectious at time of arrival into Miami-Dade County, resulting in an MLE of *R*_0_ of 0.45–0.8.

We further used this formula to address the probability of observing 3 distinct genetic clusters (F1, F2 and F3) representing 3 introduction events in a sample of 27 ZIKV genomes from Miami-Dade County. We approached this by sampling introduction events until we accumulated 241 local cases according to *L*, arriving at *N* introduction events with case counts (*j*_1_, *j*_2_,… *j*_*N*_). We then sampled 27 cases *without replacement* from (*j*_1_, *j*_2_,… *j*_*N*_) following a hypergeometric distribution and recorded the number of distinct clusters drawn in the sample. We found that higher values of *R*_0_ resulted in fewer distinct clusters within the sample of 27 genomes (Supplementary Fig. 4d). We additionally calculated the likelihood of sampling 3 distinct genetic clusters in 27 genomes (Supplementary Fig. 4e), finding an MLE estimate of *R*_0_ of 0.7–0.9. Additionally, as a sensitivity analysis we modelled a preferential sampling process in which larger clusters are more likely to be drawn from than smaller clusters. Here, we used a parameter *α* that enriches the hypergeometric distribution following (*j*_1_^*α*^, *j*_2_^*α*^,… *j*_*N*_^*α*^). In this case, we found an MLE estimate of *R*_0_ of 0.5–0.9.

Using the overlap of estimates of *R*_0_ from local case counts (0.35–0.8) and genetic clusters (0.5–0.9), we arrived at a 95% uncertainty range of *R*_0_ of 0.5–0.8.

We additionally perform birth-death stochastic simulations assuming a serial interval with mean 20 days^12^. We record the number of stochastic simulations still persisting after a particular number of days for different values of *R*_0_ (Fig. 2c).

### Incidence and attack rates

Weekly suspected and confirmed ZIKV case counts from countries and territories within the Americas with local transmission (January 1 to September 18, 2016) were obtained from the Pan American Health Organization (PAHO)^68^. In most cases, the weekly case numbers per country were only reported in bar graphs, therefore we used WebPlotDigitizer v3.10 (http://arohatgi.info/WebPlotDigitizer) to estimate the numbers. Country and territory total population sizes to calculate weekly and monthly ZIKV incidence rates were also obtained from PAHO^69^. Incidence rates calculated from countries and territories in the Americas during January to June, 2016 (based on the earliest introduction time estimates until the first known cases) were used as an estimate for infection likelihood to investigate sources of ZIKV introductions.

To account for potential reporting biases with incidence rates, we also used ZIKV attack rates (*i.e.*, proportion infected before epidemic burnout) to estimate infection likelihood. Attack rates were calculated using a susceptible–infected– recovered (SIR) transmission model derived from seroprevalence studies and environmental factors as described^70^. Using attack rates as an estimate of infection likelihood, we predict that ∼60% of the infected travelers entering Miami came from the Caribbean (Supplementary 7b), which is in agreement with our methods using incidence rates of ∼60-70% (Fig. 3c). A list of countries and territories used in these analyses can be found in Supplementary File 2.

### Airline and cruise ship traffic

To investigate whether the transmission of ZIKV in Florida coincides with travel patterns from ZIKV endemic regions, we obtained the number of passengers arriving at airports in Florida via commercial air travel. We collated flight data from countries and territories in the Americas with local ZIKV transmission between January and June, 2016 (based on the earliest introduction time estimates until the first known cases, Supplementary File 2), arriving at all commercial airports in Florida. The data were obtained from the International Air Transportation Association, which collects data on an estimated 90% of all passenger trips worldwide. Nelson et al.^26^ previously reported flight data from 33 countries with ZIKV transmission entering major United States airports during October 2014 through September 2015, which we used to assess the potential for ZIKV introductions outside of Florida.

Schedules for cruise ships visiting Miami, Port Canaveral, Port Everglades, Fort Lauderdale, Key West, Jacksonville (all in Florida), Houston, Galveston (both in Texas), Charleston (South Carolina) and New Orleans (Louisiana) ports in the year 2016 were collated from www.cruisett.com and confirmed by cross-referencing ship logs reported by Port of Miami and reported ship schedules from www.miamidade.gov/portmiami/. Scheduled cruise ship capacities were extracted from www.cruisemapper.com. Every country/territory with ZIKV transmission visited by a cruise ship 10 days (the approximate mean time to ZIKV clearance in human blood [*i.e.*, the infectious period])^71^ prior to arrival was counted as contributing the ship’s capacity worth of passengers to Miami to the month of arrival (Supplementary File 2).

### Expected number of travelers infected with Zika virus

We estimated the expected number of travelers entering Miami who were infected with ZIKV (λ) by using the total travel capacity (*C*) and the likelihood of ZIKV infection (infections (*I*) per person (*N*)) from each country/territory (*i*):

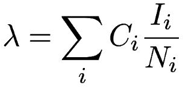

We summed the number of expected infected travelers from each country/territory with ZIKV transmission by region and travel method (flights or cruises). We reported the data as the relative proportion of infected travelers from each region (Fig. 3c, Supplementary Fig. 7a) and as the absolute number of infected travelers (Fig. 3d, Supplementary Fig. 7b, Supplementary File 2).

## Supplementary Data

**Supplementary Figure 1.**
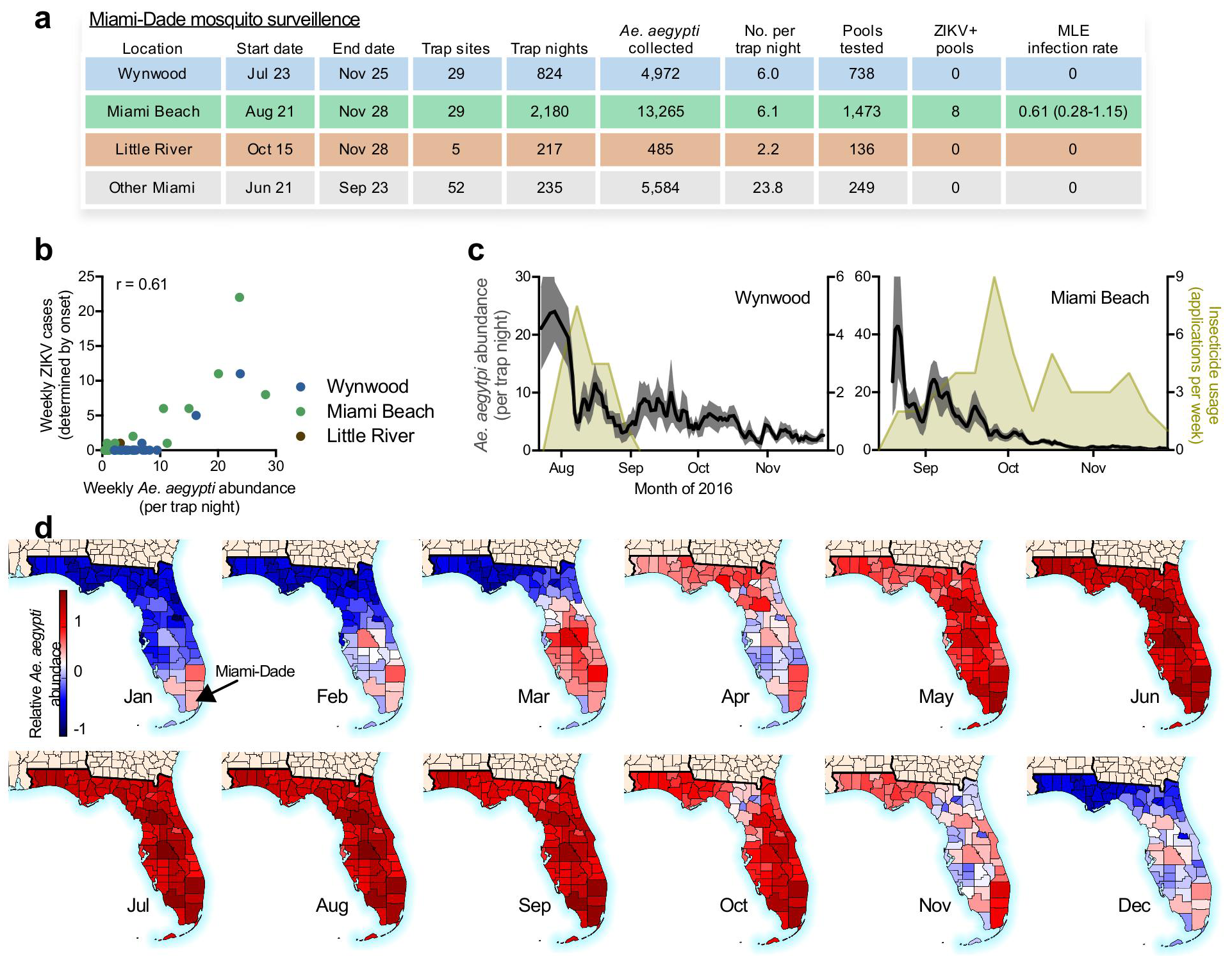
Miami-Dade mosquito surveillance and relative *Aedes aegypti* abundance. (**a**) Mosquito surveillance data reported from June 21 to November 28, 2016 was used to evaluate the risk of ZIKV infection from mosquito borne transmission in Miami. A total of 24,306 *Ae.aegypti* and 45 *Ae. albopictus* were collected. Trap nights are the total number of times each trap site was used and the trap locations are shown in Fig. 1d (some “Other Miami” trap sites are located outside of mapped region). Up to 50 mosquitoes of the same species and trap night were pooled together for ZIKV RNA testing. The infection rates were calculated using a maximum likelihood estimate (MLE). None of the *Ae. albopictus* pools contained ZIKV RNA. (**b**) The number of weekly ZIKV cases (based on symptoms onset) was correlated with mean *Ae. aegypti* abundance per trap night determined from the same week and zone (Spearman r = 0.61). This suggests that when the virus is present, mosquito abundance numbers alone could be used to target control efforts. (**c**) Insecticide usage, including truck and aerial adulticides and larvacides, by the Miami-Dade Mosquito Control in Wynwood (left) and Miami Beach (right) was overlaid with *Ae. aegypti* abundance per trap night to demonstrate that intense usage of insecticides may have helped to reduce local mosquito populations. (**d**) Relative *Ae. aegypti* abundance for each Florida county and month was estimated using a multivariate regression model, demonstrating spatial and temporal heterogeneity for the risk of ZIKV infection.

**Supplementary Figure 2.**
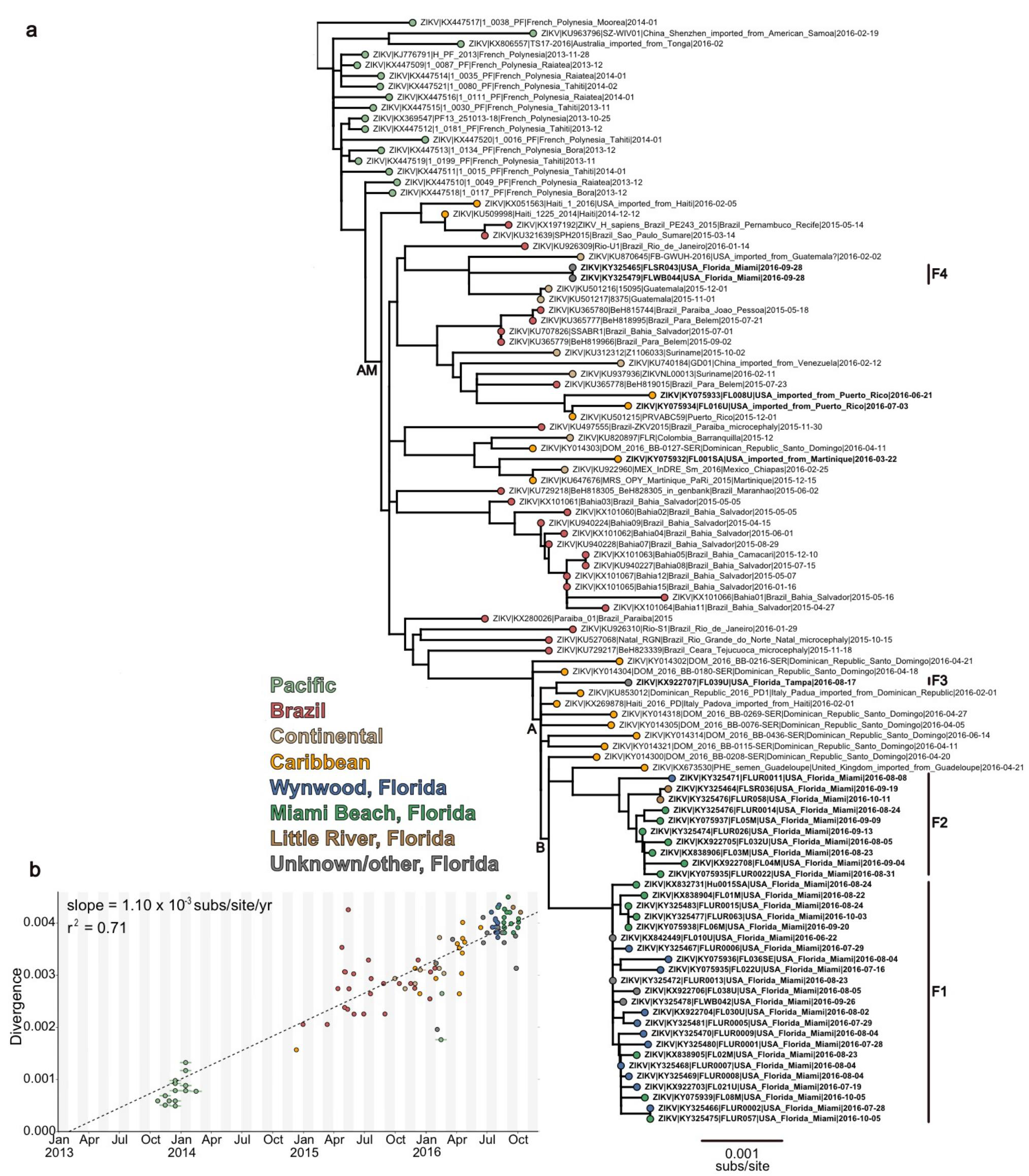
Maximum likelihood tree and root-to-tip regression of Zika virus genomes from Pacific islands and the epidemic in Americas. (**a**) Maximum likelihood tree of publicly available ZIKV sequences and sequences generated in this study (n=104). tips are coloured by location, labels in bold indicate sequences generated in this study, Florida clusters F1-F4 are indicated by vertical lines to the right of the tree. (**b**) Linear regression of sample tip dates against divergence from root based on sequences with known collection dates estimates an evolutionary rate for the ZIKV phylogeny of 1.10×10^−3^ nucleotide substitutions/site/year (subs/site/yr). This is consistent with BEAST analyses using a strict molecular clock and a Bayesian skyline tree prior, the best-performing combination of clock and demographic model according to marginal likelihood estimates (Supplementary Table 3), which estimated an evolutionary rate of 1.12×10 ^-3^ (95% highest posterior density: 0.96 - 1.27×10^−3^) subs/site/yr (Supplementary Table 1). These values are in agreement with previous estimates calculated based on ZIKV genomes from Brazil^5^.

**Supplementary Figure 3.**
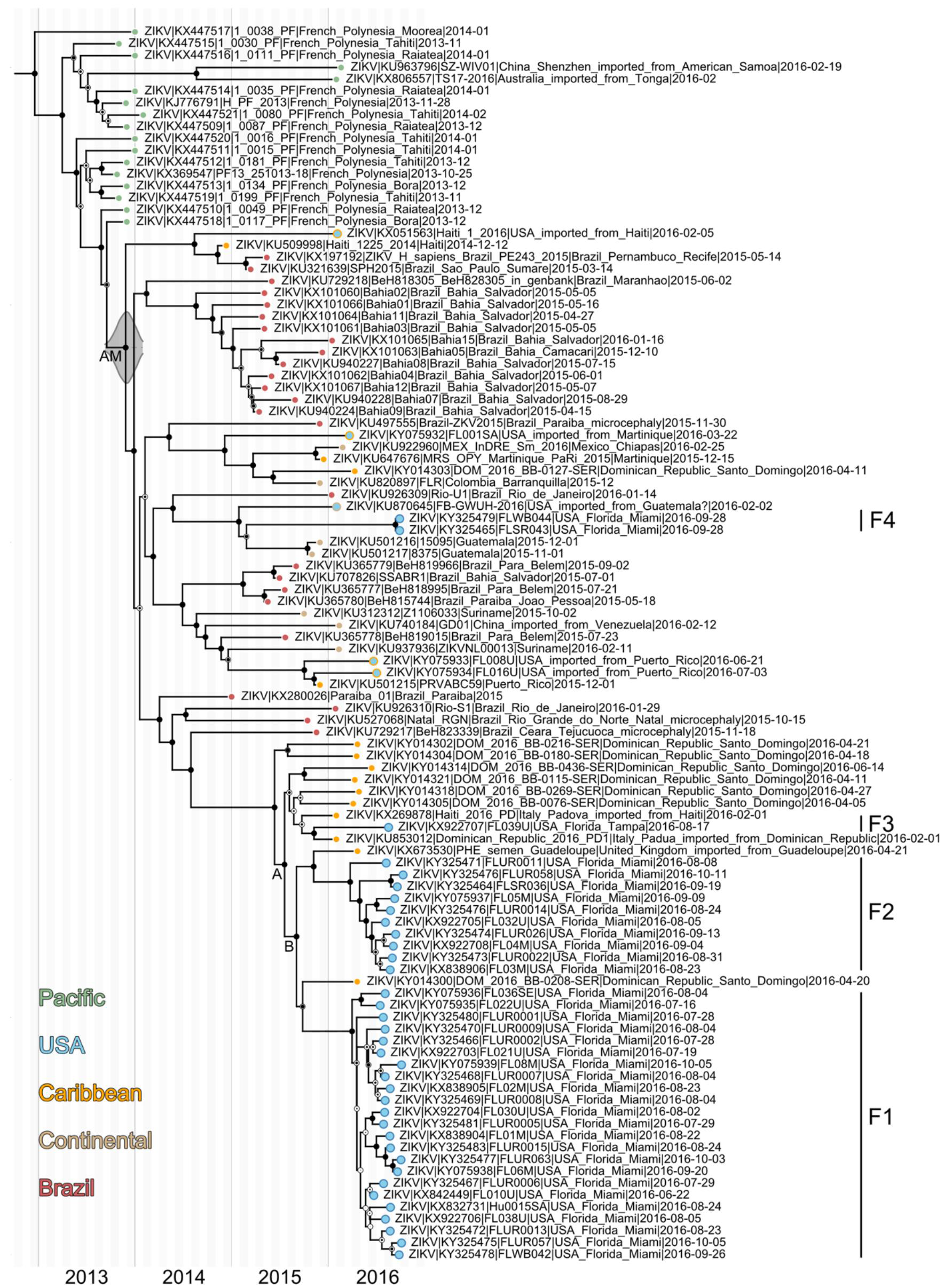
Molecular clock dating of Zika virus clades. Maximum clade credibility (MCC) tree of ZIKV genomes collected from Pacific islands and the epidemic in Americas (n=104). Circles at the tips are colored based on origin location. Clade posterior probabilities are indicated with black circles inside white circles with posterior probability of 1 filling the entire circle black. The grey violin plot indicates the 95% highest posterior density (HPD) interval for the tMRCA of the American epidemic. We estimated that the tMRCA for the ongoing epidemic in the Americas occurred during October, 2013 (node AM, Extended Table 1, 95% HPD: August, 2013-January, 2014), which is consistent with previous analysis based on ZIKV genomes from Brazil^5^.

**Supplementary Figure 4.**
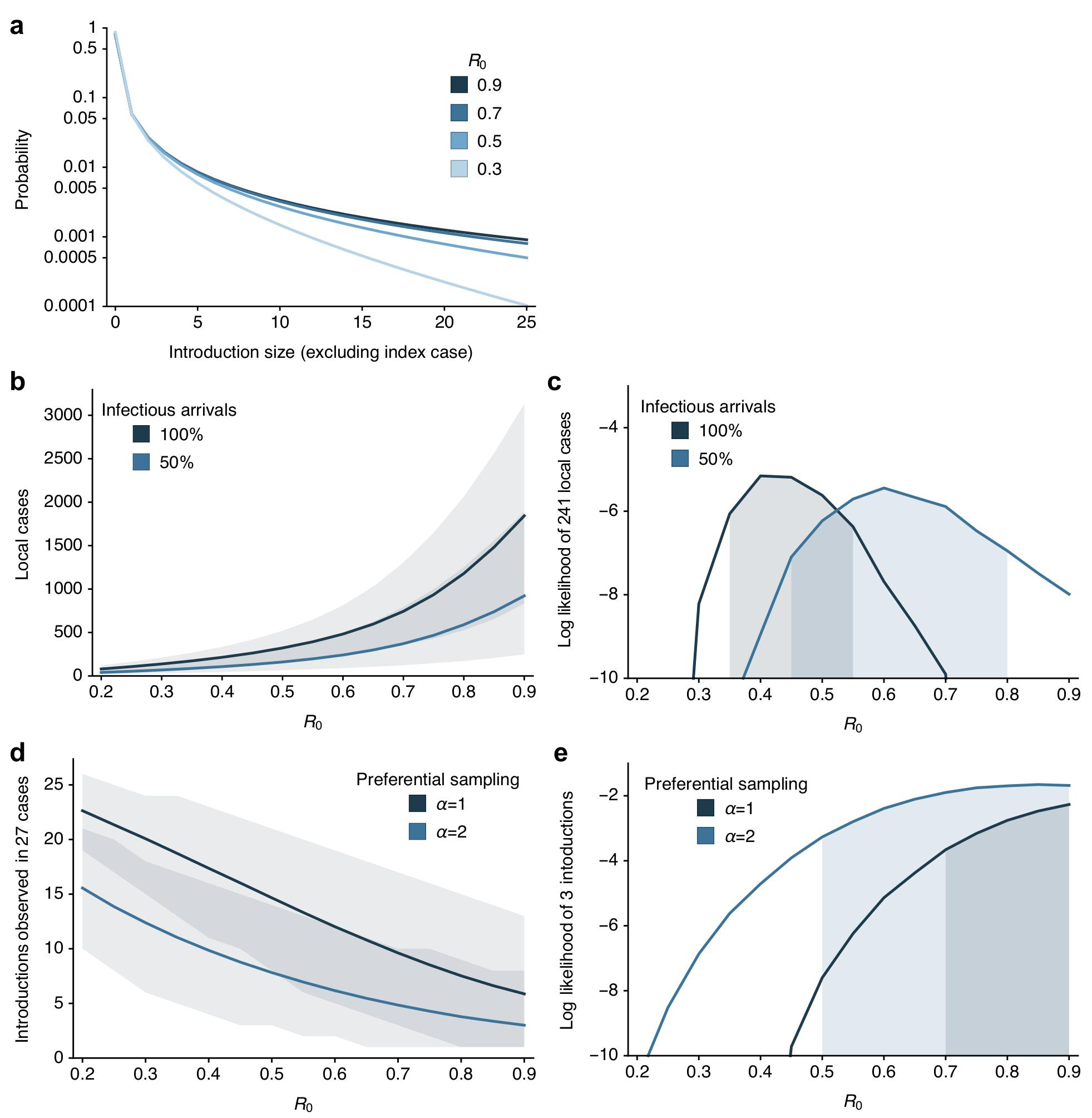
Estimation of basic reproductive number and number of introductions. (**a**) Probability distribution of estimated total number of cases caused by a single introduction (excluding the index case) for different values of *R*_0_. (**b**) Mean and 95% CI for total number of local cases caused by 320 introduction events (i*.e.*, travel-associated cases diagnosed in Miami-Dade County) for different values of *R*_0_ and for different assumptions of proportion of infectious travelers. (**c**) Log likelihood of observing 241 local cases with 320 introduction events for different values of *R*_0_ along with 95% maximum likelihood estimate (MLE) bounds on *R*_0_. (**d**) Mean and 95% uncertainty interval for total number of distinct phylogenetic clusters observed in 27 sequenced ZIKV genomes for different values of *R*_0_ and for different assumptions of sampling bias, from *α*=1 (no sampling bias) to *α*=2 (skewed toward preferentially sampling larger clusters). (**e**) Log likelihood of observing 3 clusters in 27 sequenced cases for different values of *R*_0_ along with 95% MLE bounds on *R*_0_.

**Supplementary Figure 5.**
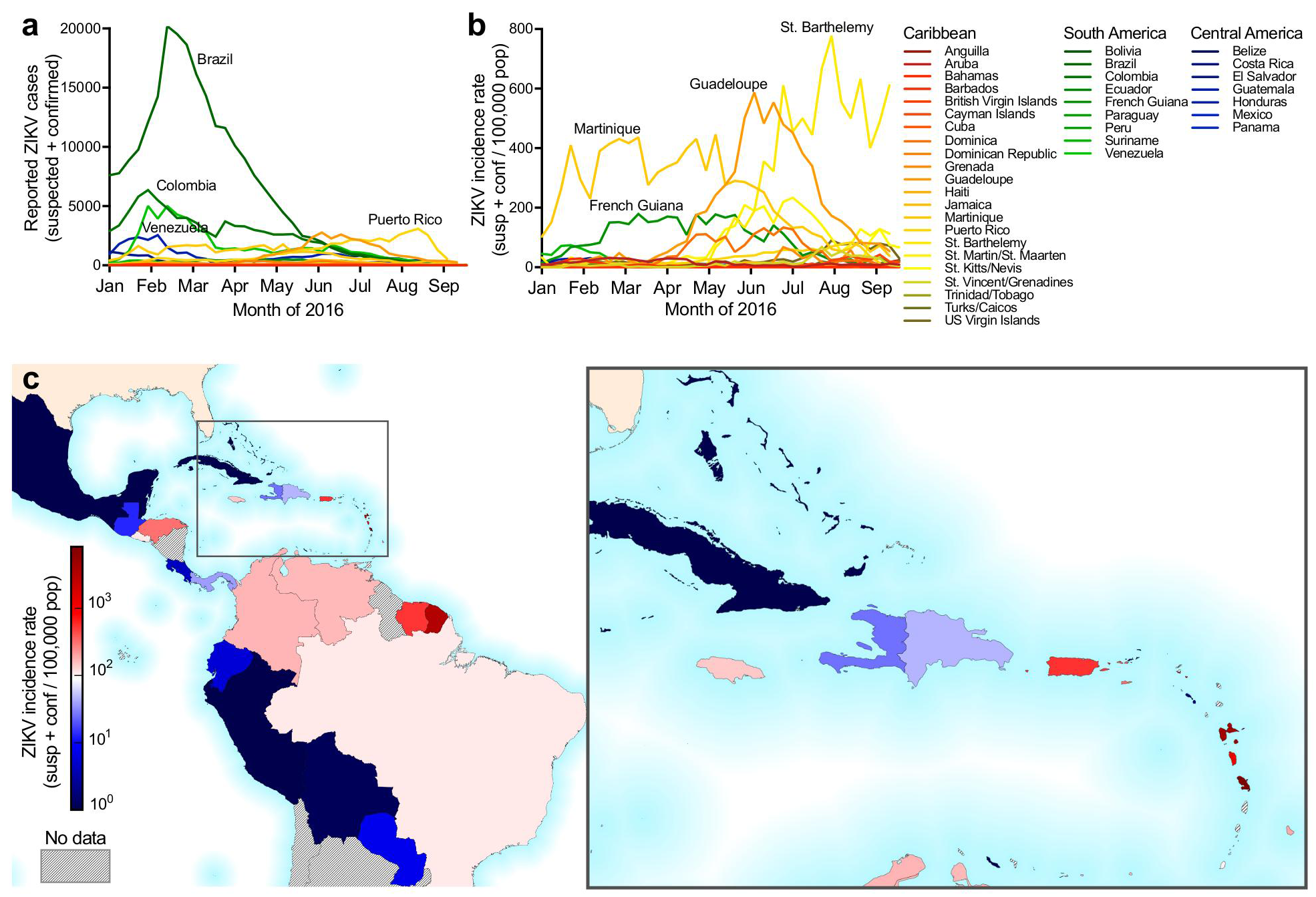
Weekly reported Zika virus case numbers and incidence rates in the Americas. (**a**) ZIKV cases (suspected and confirmed) and (**b**) incidence rates (normalized per 100,000 population) are shown for each country or territory with available data per epidemiological week from January 1 to September 18, 2016. (**c**) Each country or territory with available data is colored by its reported ZIKV incidence rate from January to June, 2016 (the time frame for analysis of ZIKV introductions into Florida).

**Supplementary Figure 6.**
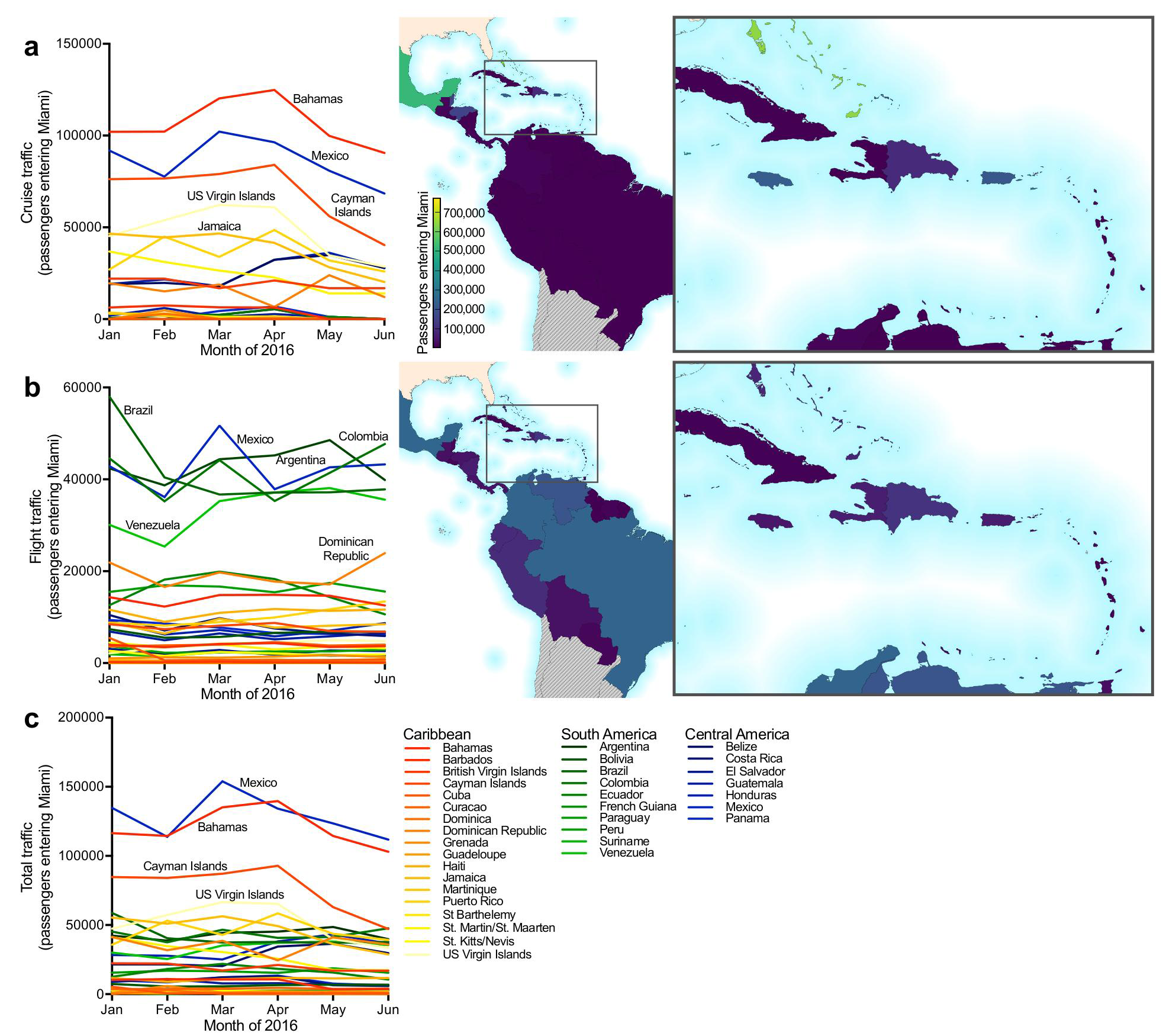
Cruise and flight traffic entering Miami from regions with Zika virus transmission. The expected number of passengers entering Miami, by either (**a**) cruises or (**b**) flights, from each country or territory in the Americas with ZIKV transmission per month (left panel). The center map and inset show the cumulative numbers of travelers entering Miami during January to June, 2016 (the time frame for analysis of ZIKV introductions into Florida) from each country or territory per method of travel. (**c**) The total traffic (*i.e.* cruises and flights) is shown entering Miami per month.

**Supplementary Figure 7.**
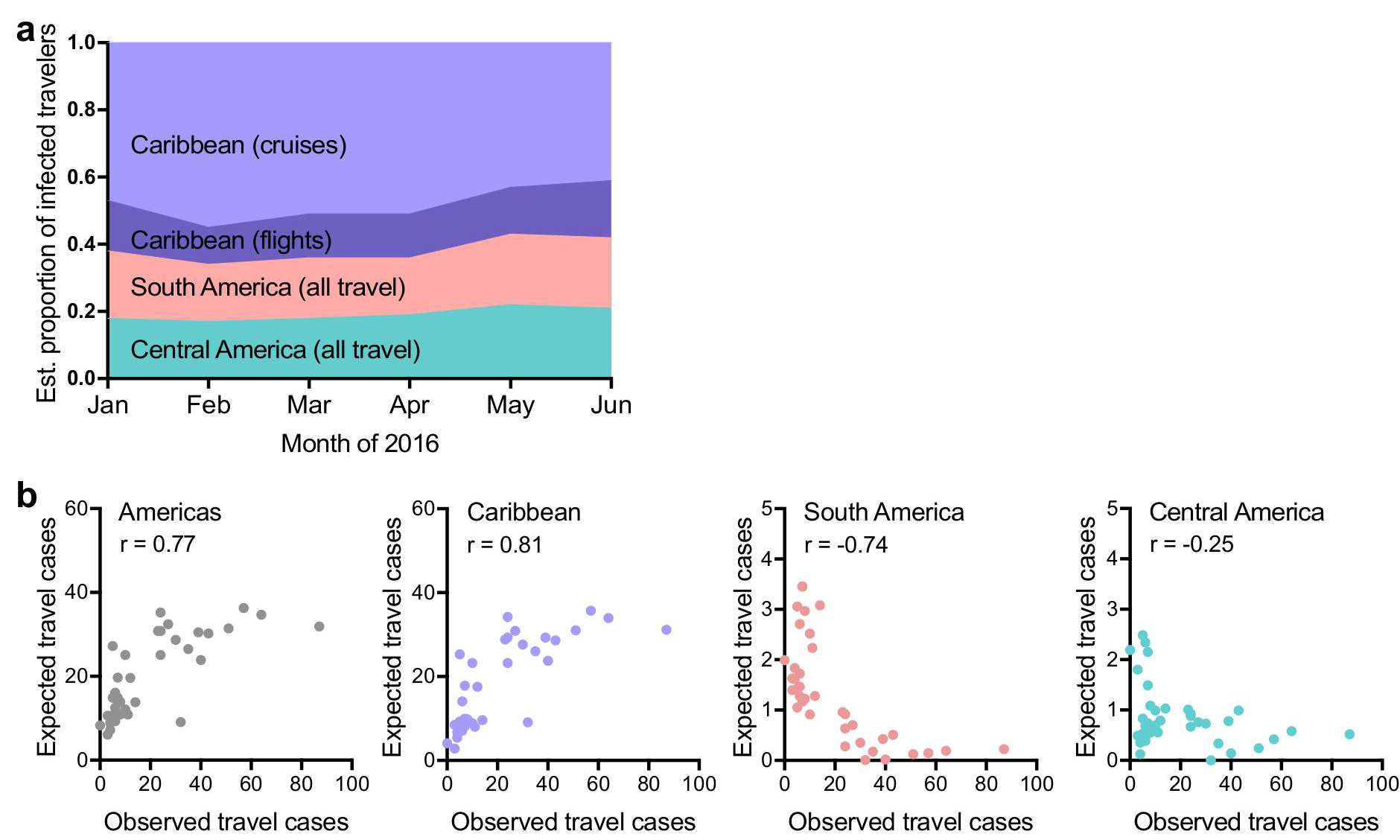
Expected number of Zika virus infected travelers from the Caribbean is correlated with the total observed number of travel-associated infections. (**a**) In order to account for potential biases in ZIKV reporting accuracies, we also estimated the proportion of infected travelers using projected ZIKV attack rates ^70^ (*i.e.* predicted proportion of population infected before epidemic burnout). About 60% of the infected travelers are expected to have arrived from the Caribbean, similar to our results using incidence rates (**Fig. 3c**). (**b**) The expected number of travel-associated ZIKV cases were estimated by the number of travelers coming into Miami from each country/territory (travel capacity) and the in-country/territory infection likelihood (incidence rate per person) per week. The expected travel cases were summed from all of the Americas (left), Caribbean (left center), South America (right center), and Central America (right) and plotted with the observed travel-associated ZIKV cases. Numbers in each plot indicate Spearman correlation coefficients. Negative Spearman r coefficients indicated a negative correlation between the number of expected and observed travel cases.

**Supplementary Figure 8.**
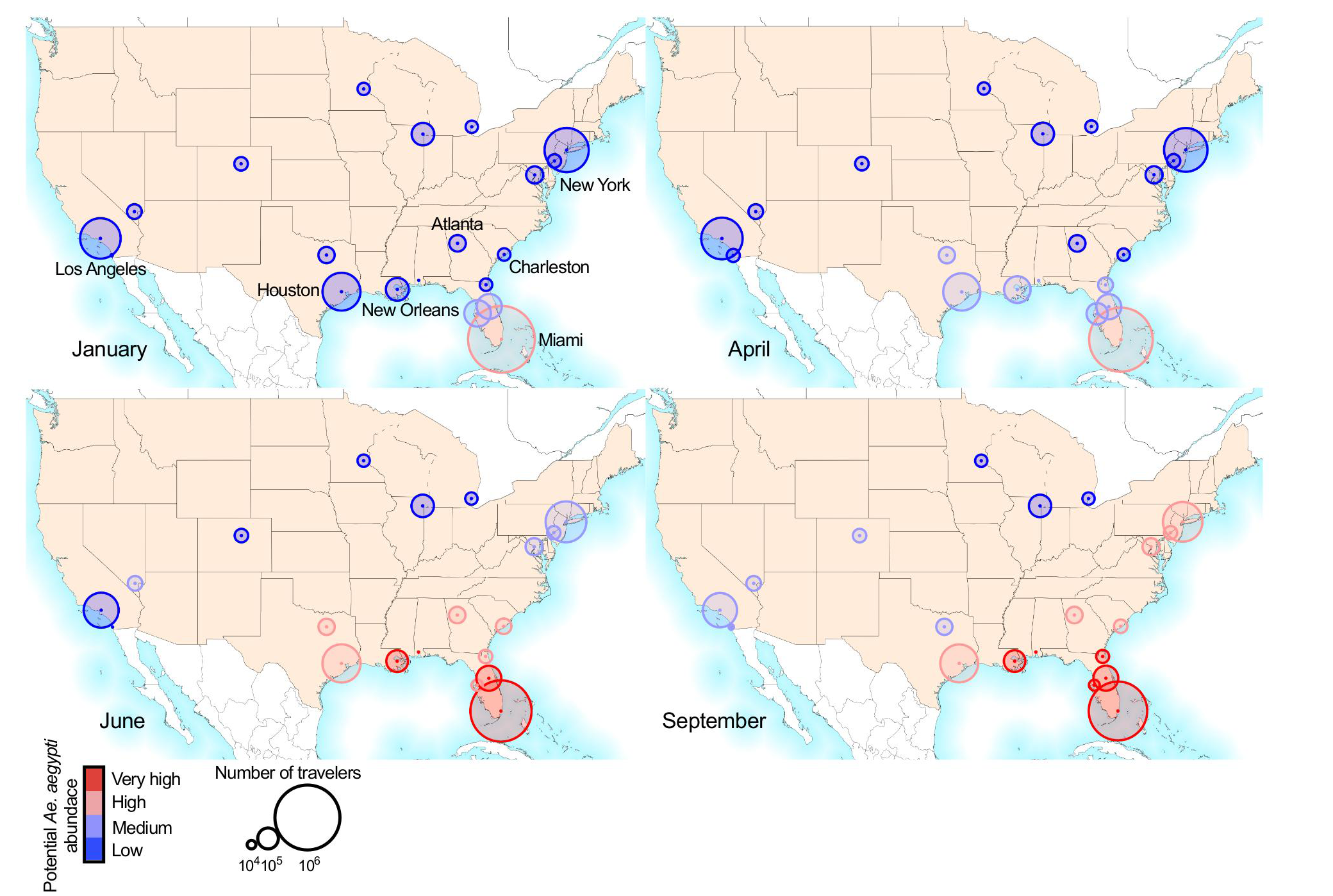
Greater early season potential for Zika virus introductions into Miami. (**a**) The monthly cruise ship and airline^26^capacity from countries/territories with ZIKV transmission for the major United States travel hubs (shown as circle diameter) with monthly potential *Ae.aegypti* abundance (circle color), as previously estimated^20^. Cruise capacities from Houston and Galveston, Texas were combined.

**Supplementary Table 1.**
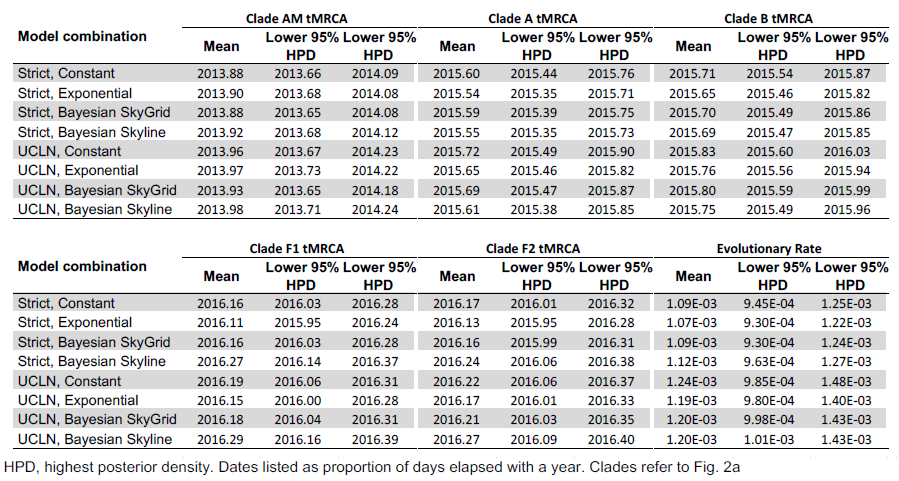
Times of the most recent common ancestor and evolutionary rates

| Model combination | Clade AM tMRCA |  |  | Clade A tMRCA |  |  | Clade B tMRCA |  |  |
| --- | --- | --- | --- | --- | --- | --- | --- | --- | --- |
|  | Mean | Lower 95% HPD | Lower 95% HPD | Mean | Lower 95% HPD | Lower 95% HPD | Mean | Lower 95% HPD | Lower 95% HPD |
| Strict, Constant | 2013.88 | 2013.66 | 2014.09 | 2015.60 | 2015.44 | 2015.76 | 2015.71 | 2015.54 | 2015.87 |
| Strict, Exponential | 2013.90 | 2013.68 | 2014.08 | 2015.54 | 2015.35 | 2015.71 | 2015.65 | 2015.46 | 2015.82 |
| Strict, Bayesian SkyGrid | 2013.88 | 2013.65 | 2014.08 | 2015.59 | 2015.39 | 2015.75 | 2015.70 | 2015.49 | 2015.86 |
| Strict, Bayesian Skyline | 2013.92 | 2013.68 | 2014.12 | 2015.55 | 2015.35 | 2015.73 | 2015.69 | 2015.47 | 2015.85 |
| UCLN, Constant | 2013.96 | 2013.67 | 2014.23 | 2015.72 | 2015.49 | 2015.90 | 2015.83 | 2015.60 | 2016.03 |
| UCLN, Exponential | 2013.97 | 2013.73 | 2014.22 | 2015.65 | 2015.46 | 2015.82 | 2015.76 | 2015.56 | 2015.94 |
| UCLN, Bayesian SkyGrid | 2013.93 | 2013.65 | 2014.18 | 2015.69 | 2015.47 | 2015.87 | 2015.80 | 2015.59 | 2015.99 |
| UCLN, Bayesian Skyline | 2013.98 | 2013.71 | 2014.24 | 2015.61 | 2015.38 | 2015.85 | 2015.75 | 2015.49 | 2015.96 |

| Model combination | Clade F1 tMRCA |  |  | Clade F2 tMRCA |  |  | Evolutionary Rate |  |  |
| --- | --- | --- | --- | --- | --- | --- | --- | --- | --- |
|  | Mean | Lower 95% HPD | Lower 95% HPD | Mean | Lower 95% HPD | Lower 95% HPD | Mean | Lower 95% HPD | Lower 95% HPD |
| Strict, Constant | 2016.16 | 2016.03 | 2016.28 | 2016.17 | 2016.01 | 2016.32 | 1.09E-03 | 9.45E-04 | 1.25E-03 |
| Strict, Exponential | 2016.11 | 2015.95 | 2016.24 | 2016.13 | 2015.95 | 2016.28 | 1.07E-03 | 9.30E-04 | 1.22E-03 |
| Strict, Bayesian SkyGrid | 2016.16 | 2016.03 | 2016.28 | 2016.16 | 2015.99 | 2016.31 | 1.09E-03 | 9.30E-04 | 1.24E-03 |
| Strict, Bayesian Skyline | 2016.27 | 2016.14 | 2016.37 | 2016.24 | 2016.06 | 2016.38 | 1.12E-03 | 9.63E-04 | 1.27E-03 |
| UCLN, Constant | 2016.19 | 2016.06 | 2016.31 | 2016.22 | 2016.06 | 2016.37 | 1.24E-03 | 9.85E-04 | 1.48E-03 |
| UCLN, Exponential | 2016.15 | 2016.00 | 2016.28 | 2016.17 | 2016.01 | 2016.33 | 1.19E-03 | 9.80E-04 | 1.40E-03 |
| UCLN, Bayesian SkyGrid | 2016.18 | 2016.04 | 2016.31 | 2016.21 | 2016.03 | 2016.35 | 1.20E-03 | 9.98E-04 | 1.43E-03 |
| UCLN, Bayesian Skyline | 2016.29 | 2016.16 | 2016.39 | 2016.27 | 2016.09 | 2016.40 | 1.20E-03 | 1.01E-03 | 1.43E-03 |

HPD, highest posterior density. Dates listed as proportion of days elapsed with a year. Clades refer to Fig. 2a

**Supplementary Table 2.**
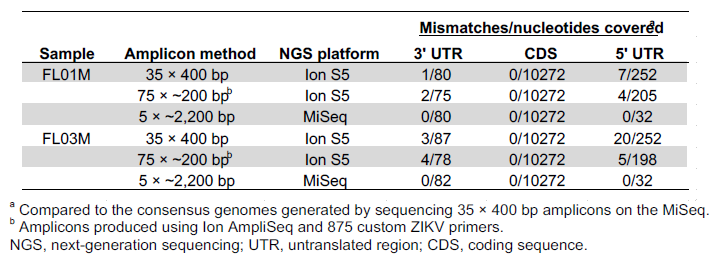
Validation of sequencing results.

**Supplementary Table 3.**
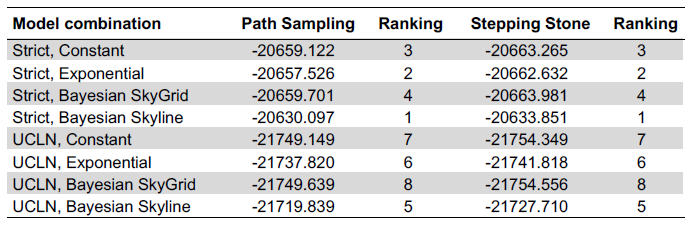
Model selection to infer time-structured phylogenies.

| Model combination | Path Sampling | Ranking | Stepping Stone | Ranking |
| --- | --- | --- | --- | --- |
| Strict, Constant | -20659.122 | 3 | -20663.265 | 3 |
| Strict, Exponential | -20657.526 | 2 | -20662.632 | 2 |
| Strict, Bayesian SkyGrid | -20659.701 | 4 | -20663.981 | 4 |
| Strict, Bayesian Skyline | -20630.097 | 1 | -20633.851 | 1 |
| UCLN, Constant | -21749.149 | 7 | -21754.349 | 7 |
| UCLN, Exponential | -21737.820 | 6 | -21741.818 | 6 |
| UCLN, Bayesian SkyGrid | -21749.639 | 8 | -21754.556 | 8 |
| UCLN, Bayesian Skyline | -21719.839 | 5 | -21727.710 | 5 |

**Supplementary File 1.** Summary of the Zika virus sequencing data produced in this study.

**Supplementary File 2.** Epidemiological data and travelers entering Miami, Florida from January to June, 2016.

**Supplementary File 3.** Primer sequences used for long-range amplicons and probe sequences for RNA Access targeted enrichment of Zika virus.

**Supplementary File 4.** Raw MAFFT codon alignment, PhyML tree, BEAST XML file, and BEAST MCC time-structured phylogeny.

